# VesSynth: Tubes Are All You Need for Robust Cross-Scale Cross-Modal 3D Vessel Segmentation

**DOI:** 10.64898/2026.04.01.715909

**Authors:** Chiara Mauri, Allison McKenzie, Cole Analoro, Emma Yeon, Rose Coviello, Jocelyn Mora, Etienne Chollet, Lucas Deden Binder, Ara Mahar, Stephanie Lin, Malak Benlahcen, Angelina Ream, Aliyah Jama, Itzel Garcia, Nam Tran, Priyanka Onta, Sariya Wood, Adam Willis, Alisha Mahmood, Greisi Sinoballa, Akram Malki, Kenton Tran, Vennela Malireddy, Nkiruka Onumajuru, Sonia Lakshmanan, Kaylee Hercules Landaverde, Rahama Sidow, David Wood, Binh Nguyen, Jiuver Hernandez, Maggie Bernier, Jayvi Hunter, Achraf Malki, Annabella Tum, Victoria Chavez, Zenera Shahu, Isabella Vasi, Abigail Visser, Zahra Ghaouta, Felicia Bond, Rithikaa Vigneshwaran, Emilia Kirkpatrick, Michelle Avalos Barbosa, Kathryn Rauh, Rogeny Herisse, Erendira Garcia Pallares, Xiangrui Zeng, Divya Varadarajan, Hui Wang, Caroline Magnain, Brian L. Edlow, Malte Hoffmann, Bruce Fischl, Yaël Balbastre

## Abstract

The cerebral vasculature is central to brain function, with alterations linked to numerous cerebrovascular and neurological disorders. Yet, no single imaging modality can capture the entire cerebral vascular network in humans. Instead, an array of techniques are sensitized to different spatial scales, while trading off resolution for coverage. Magnetic Resonance Imaging (MRI) typically resolves only large pial vessels, while high-resolution microscopy allows micrometer-scale vessels to be mapped over limited spatial extents. These techniques must therefore be combined to obtain a complete mapping of the cerebral angioarchitecture, which underscores the need for automatic, cross-modal vessel segmentation. Here, we introduce VesSynth, a flexible vessel segmentation framework that achieves state-of-the-art accuracy across multiple modalities and spatial resolutions (MR, optical and X-ray imaging), despite being trained entirely on synthetic data. By enabling consistent vascular mapping across scales, this framework paves the way to comprehensive investigation of cerebrovascular organization and its role in health and disease.

---

The cerebral vasculature is a fundamental component of brain health. Its alterations are involved in a wide spectrum of disorders, including cerebrovascular, neurodegenerative, and psychiatric conditions, positioning cerebrovascular health as a key factor across brain diseases. Among cerebrovascular conditions, cerebral small vessel disease (CSVD) is a major subtype affecting small brain vessels [1]; it encompasses conditions such as cerebral amyloid angiopathy, where pathogenic proteins deposit along vessel walls and contribute to impaired brain waste clearance in perivascular pathways and cognitive decline [2, 3, 4]. Overall, CSVD represents a major public health burden, contributing substantially to stroke [5, 6] and dementia [6, 7], often co-occuring with Alzheimer’s disease [5], and yielding increased risk of late-life depression, [8, 9, 10], marked by increased cognitive impairment and a poorer response to treatment compared to depression earlier in life [11]. More broadly, growing evidence highlights the role of vascular dysfunction beyond purely vascular diseases. Vascular alterations– including protein accumulation along vessels, which impairs waste clearance mechanisms, blood–brain barrier disruption, impaired cerebral blood flow, and dysregulated angiogenesis– are implicated in a wide range of disorders, from Alzheimer’s disease [12, 13, 14, 15], to stroke [15, 16], Parkinson’s disease [17], multiple sclerosis [17], and cancer [15, 16].

Given the growing body of evidence implicating cerebrovascular health as a critical determinant across multiple brain disorders, there is a pressing need for robust methodologies to characterize vascular morphometry and how it is altered in neuropathological conditions. In small mammals, novel imaging methods have led to the mapping of the entire cerebral vascular network [18]. However, the human brain is four orders of magnitude larger than the mouse brain, and its cerebral vessels span a hierarchy from millimeter-scale pial arteries and veins to capillaries with diameters of only a few micrometers. Studying the vasculature across such different spatial scales requires a combination of imaging modalities, each leveraging a different contrast mechanism. Magnetic Resonance Angiography (MRA) enables *in vivo* visualization of arterial structures with high contrast. While traditionally limited to resolving only the largest pial vessels, recent advances with ultra-high field have pushed MRA into the mesoscopic regime (*c*. 150 µm) [19].

Similarly, T2*-weighted contrast mechanisms, such as those used in susceptibility-weighted imaging, permit *in vivo* visualization of venous structures, with state-of-the-art techniques now enabling spatial resolutions on the order of 300 µm at ultrahigh field [20]. *Ex vivo* MRI further extends imaging capabilities by providing mesoscopic resolution (*c*. 100 µm) with whole-brain coverage [21], enabling visualization of some penetrating arterioles and venules, and most medullary veins, although with lower signal-to-noise ratio and less specific vessel contrast. At the highest resolution, detailed knowledge of vascular anatomy down to the capillary level comes primarily from microscopy techniques such as light-sheet fluorescence microscopy and electron microscopy, which provide micrometer- and nanometer-scale detail but are limited in spatial extent [22, 23].

Traditionally, segmentation methods have utilized known geometric features of the vessels *via* Hessian-based filters [24], morphological operations [25], or region-growing techniques [26]. They are computationally efficient but degrade in the presence of noise, artifacts, or unexpected morphology [27, 28, 29]. Deep learning has since revolutionized semantic segmentation, particularly through the use of convolutional neural networks, which have significantly improved performance in challenging conditions [30, 31]. However, their effectiveness remains strongly dependent on the amount and quality of manually annotated data used for training, highlighting a key limitation in settings with scarce and difficult-to-obtain labels. Most deep learning methods for vessel segmentation are tailored to a specific imaging modality and spatial scale [31, 30, 32, 33], and hardly generalize across domains. Recent work in machine learning has focused on foundation models, in which data size, model size, and task diversity are massively scaled up [34]. Foundation models improve generalization to new domains and tasks, either with or without fine-tuning. While most foundation models target very general domains, such as object segmentation [35], there is a trend towards training foundation models dedicated to more specialized applications [36]. In this context, two novel approaches have been developed with the aim of improving cross-modal generalization of vascular segmentation: Holroyd and colleagues trained a foundation model across multiple modalities with subsequent fine-tuning on new domains [37], while Wittmann and colleagues complemented this approach with synthetic training data and domain randomization, which allowed them to improve performance in zero-shot settings [38]. Despite curating multiple open datasets, these models were trained on only *c*. 400 individual images, the majority of which are low-resolution MR or CT images. Despite some of these images being large, this vastly limits the variety of vascular anatomies that models learn from. Furthermore, expert annotations are often incomplete due to the fine and fractal structure of the vasculature, which ensures that for any given imaging system, some vascular segments lie at its resolution limit. Consequently, these models obtain much higher performance on modalities and resolutions that are present in their training data than on entirely new domains [38].

To address the scarcity of high-quality manual annotations for vessels and facilitate rapid adaptation to new imaging modalities, we introduce *VesSynth*, a flexible and robust framework for 3D vessel segmentation trained entirely on synthetic data. The framework combines spline-based synthesis and domain randomization to train robust segmentation models without manual annotations [39], while facilitating rapid adaptation to new imaging modalities [40, 41]. The use of synthetic training data has been widely shown to enable state-of-the-art generalization in medical image analysis tasks [41, 42, 43, 44]. In the context of vessel segmentation, the synthetic-data paradigm offers several key advantages: (1) access to effectively unlimited, perfectly paired training data, avoiding the limitations of imperfect manual annotations; (2) strong generalization across datasets within the same modality, regardless of scanner or acquisition protocol [42]; and (3) high flexibility, as the synthesis pipeline, while modality-specific, can be easily extended to other imaging modalities. Here, we employ the framework to develop automatic vessel segmentation across imaging modalities that span multiple spatial scales of the human brain: **Time-of-flight Magnetic Resonance Angiography** (MRA-TOF; 150–800 µm/voxel), ***Ex vivo* MRI** (100–150 µm/voxel), **Hierarchical Phase-Contrast Tomography** (HiP-CT; 10–50 µm/voxel) and **Optical Coherence Tomography** (OCT; 5–20 µm/voxel). VesSynth achieves state-of-art performances across this diverse set of imaging modalities and is publicly available at https://github.com/chiara-mauri/VesSynth.

## Results

### A carefully labeled validation dataset across scales and modalities

For validation and testing of the segmentation method, we acquired and/or curated images from four imaging modalities:

- **Time-of-flight Magnetic Resonance Angiography (MRA-TOF)** is an *in vivo* technique for visualizing blood vessels, primarily arteries, without the need for contrast agents. It exploits the difference in signal between stationary tissue that has been saturated by radio-frequency pulses and unsaturated blood flowing into an imaging slab, resulting in a strong vascular signal (“inflow effect”). Spatial resolution ranges from millimeter to sub-millimeter, with advanced protocols achieving higher resolutions (*c*. 150 µm) [19].
- ***Ex vivo* MRI**provides whole-brain imaging at mesoscopic resolution (100–150 µm) resolution, allowing visualization of penetrating arterioles, venules, and medullary veins— structures typically beyond the reach of *in vivo* MRI. Automatic vessel segmentation in *ex vivo* MRI can enable investigation of morphology and inter-subject variability in vascular architecture at mesoscopic scales. However, this modality poses challenges due to its inherently low signal-to-noise ratio and the limited contrast between vessels and surrounding parenchyma.
- **Hierarchical Phase-Contrast Tomography (HiP-CT)** is an X-ray phase propagation technique that enables ultra-high resolution 3D imaging of tissue without physical sectioning [45]. HiP-CT imaging is performed hierarchically: at its coarsest scale, it provides whole-organ scans at 10–30 µm isotropic voxel size. Within the same sample, selected volumes of interest can then be imaged at much finer resolution, down to 1 µm voxel size, without physical sectioning. This hierarchical approach bridges the gap between macroscopic imaging (whole-organ morphology) and microscopic detail.
- **Optical Coherence Tomography (OCT)** is an optical technique that images tissue blocks by quantifying the intensity of back-scattered light from the tissue. It provides high spa-tial resolution (*c*. 5–20 µm) and can resolve small arterioles and venules with diameters down to 20 µm [27]. The field of view typically spans several hundred micrometers in depth and, when used in combination with a tissue slicer, can extend to a multiple centimeters through-plane. Notably, OCT limits tissue distortion by acquiring images directly from the block face prior to sectioning.

We then manually annotated vessels on small volumetric patches (64^3^ voxels) extracted from these images, as shown in Fig. 1. The small patch size was chosen to facilitate exhaustive annotation, as vessels are ubiquitous throughout the tissue and span a wide range of spatial scales, including near the edge of visibility. Every patch was labeled across multiple passes by multiples operators, ensuring the highest possible level of accuracy. These annotations were not used for training, as the segmentation model was trained exclusively on synthetic data, but served for model selection and independent evaluation. The MRA-TOF dataset was further divided into a lower-resolution subset (test set A) and a higher-resolution subset (test set B) to assess performance across imaging conditions.

**Figure 1:**
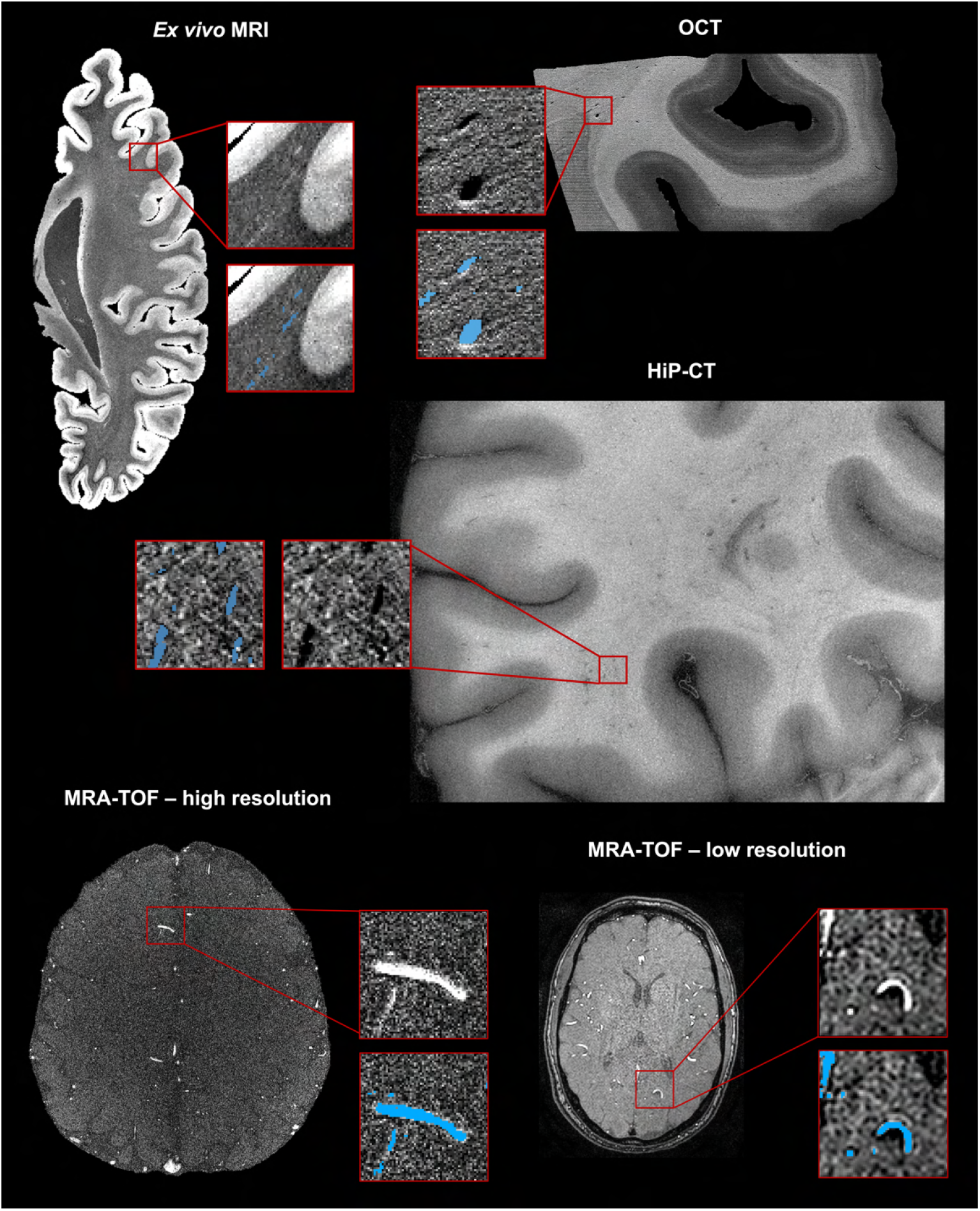
Examples of real data for each modality, including extracted 64^3^-voxel patches with corresponding manual vessel labels, used for validation and testing. In the last row, the MRA-TOF image on the left has 0.16 mm isotropic resolution, while the one on the right has resolution 0.47 × 0.47 × 0.80 mm.

### Spline-based synthesis can flexibly be tuned to target a diverse range of image modalities

For each modality we generated synthetic vascular trees using B-splines and paired them with modality-specific synthetic intensity images (Fig. 2). Both vessel geometry and intensity synthesis were intentionally extended beyond realistic appearance to expose the network to a broad range of structural and contrast variability. The generated synthetic data were used to train modality-specific vessel segmentation models using a U-Net architecture [46]. Details of the synthesis pipeline and training procedure are provided in the Methods section.

**Figure 2:**
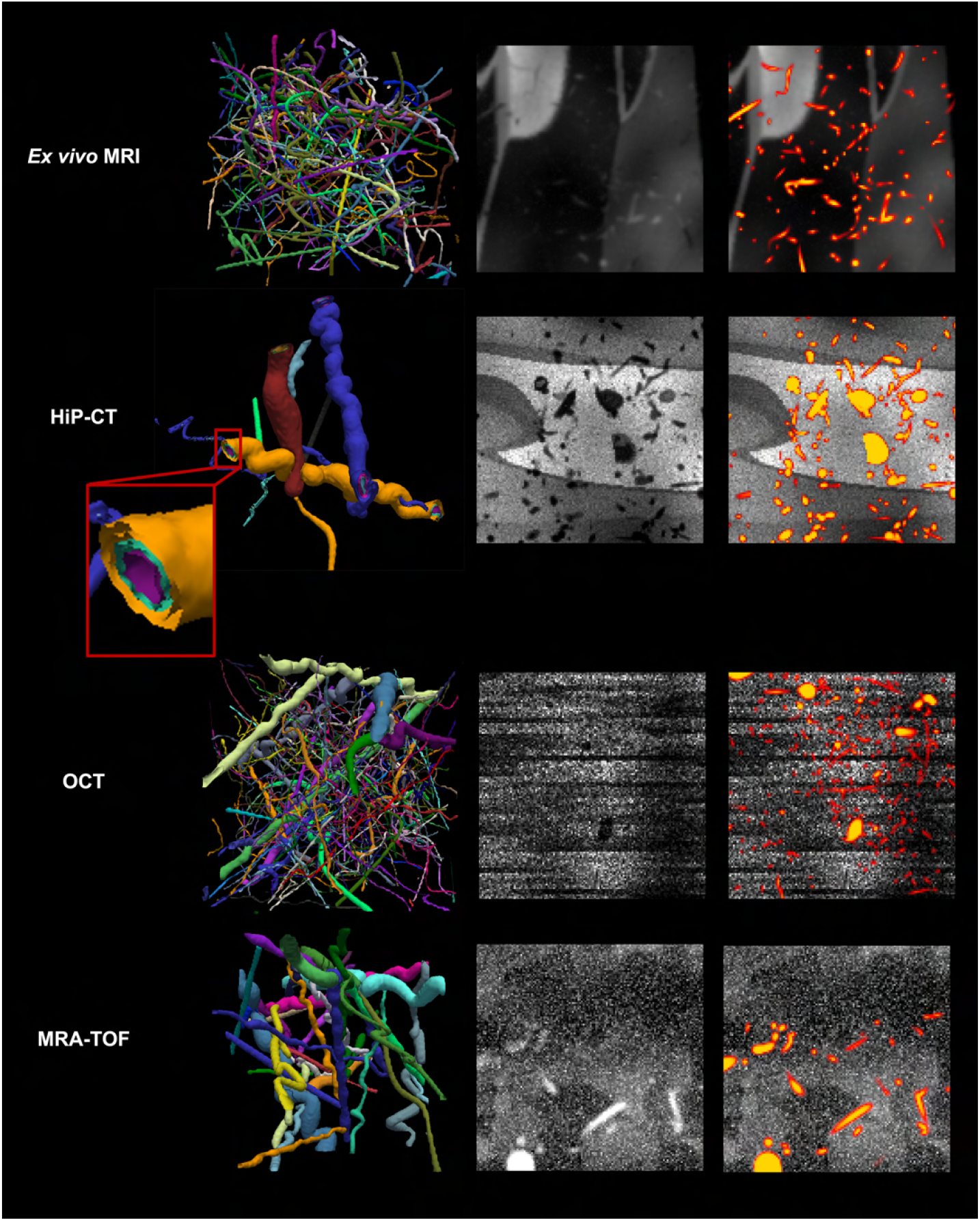
Left column: 3D rendering of synthetic vascular trees, where each vessel is assigned a unique ID and visualized with a distinct color. The HiP-CT panel shows a close-up of the three vascular compartments. Middle column: 2D view of corresponding synthetic intensity images. Right column: synthetic intensity images with the true vessel probability map overlaid. This figure illustrates a single example from the stochastic synthesis process and is therefore not exhaustive.

### Validation metrics and baselines

Segmentation accuracy was evaluated on a test set of real, manually labeled 64^3^-voxel patches using several metrics: *Dice score* (DSC) [47], *soft* DSC, *corrected* DSC; area under the receiver operating characteristic curve (AUC); *corrected* true positive rate (TPR); and *corrected* false discovery rate (FDR). We compared VesSynth against the following benchmarks on the same test sets: (1) a supervised U-Net that we trained *on real data* for each modality; (2) Frangi filters, a modality-agnostic method for tubular structure segmentation [24]; (3) VesselFM [38], a state-of-the-art deep learning foundation model for vessel segmentation, trained on a mixture of real and synthetic data; and (4) for MRA-TOF only, VesselBoost [32], a state-of-the-art deep learning model for MRA-TOF vessel segmentation. Further details on evaluation metrics and benchmarks are provided in the Methods section.

### VesSynth either matches or exceeds the performance of supervised and foundation baselines across all modalities

Results are reported in Fig. 3 (corrected DSC) and Supplementary Table 3 (all other metrics). For *ex vivo* MRI, HiP-CT, and OCT, the proposed method achieves the highest corrected DSC across all benchmarks (0.83–0.89), demonstrating strong performance across modalities. For MRA-TOF, on test set A (lower resolution), the supervised U-Net trained on real data achieves the highest performance, closely followed by the proposed method (mean corrected DSC 0.95 vs. 0.94). On test set B instead the proposed method and VesselFM achieve the best results (mean corrected DSC 0.92 for both). VesselFM achieves strong performance on HiP-CT and MRA-TOF (mean corrected DSC > 0.90), but more modest performance on *ex vivo* MRI and OCT (0.45 and 0.59, respectively). This likely reflects that, despite training on a mixture of real and synthetic data, the model performs best on modalities represented in its real training set (HiP-CT and MRA-TOF, see Methods), and less well on others. We note that VesselFM is contrast-agnostic, whereas the other models are tuned to specific modalities. Frangi filters perform poorly on HiP-CT and OCT (0.31 and 0.45 respectively), and modestly on ex vivo MRI and MRA-TOF higher resolution (0.61 and 0.63), likely due to their sensitivity to noise, reaching higher accuracy only on MRA-TOF test set A (0.89). The supervised U-Net achieves mean corrected DSC values ranging from 0.71 to 0.95 across modalities. Aggressive data augmentation during training, including geometric transformations and intensity artifacts, was essential to achieve strong performance (results not shown), highlighting the importance of exposing the model to increased variability. The strong performance of the supervised U-Net on MRA-TOF test set A, where it is the top-performing method, likely reflects both the larger number of training patches available for this modality (supplementary Table 2a) and the higher vessel contrast, which facilitates more accurate manual annotations. Its drop in performance from MRA-TOF test set A (0.95) to test set B (0.86) likely reflects the similarity of test set A to its training data, whereas test set B presents higher resolution and field strength (Supplementary Table 2b), leading to reduced accuracy. In contrast, the synthetic-data approach of the proposed method yields more robust generalization to these unseen conditions. Across both MRA-TOF test sets, the proposed method consistently outperforms Frangi filters and VesselBoost.

**Figure 3:**
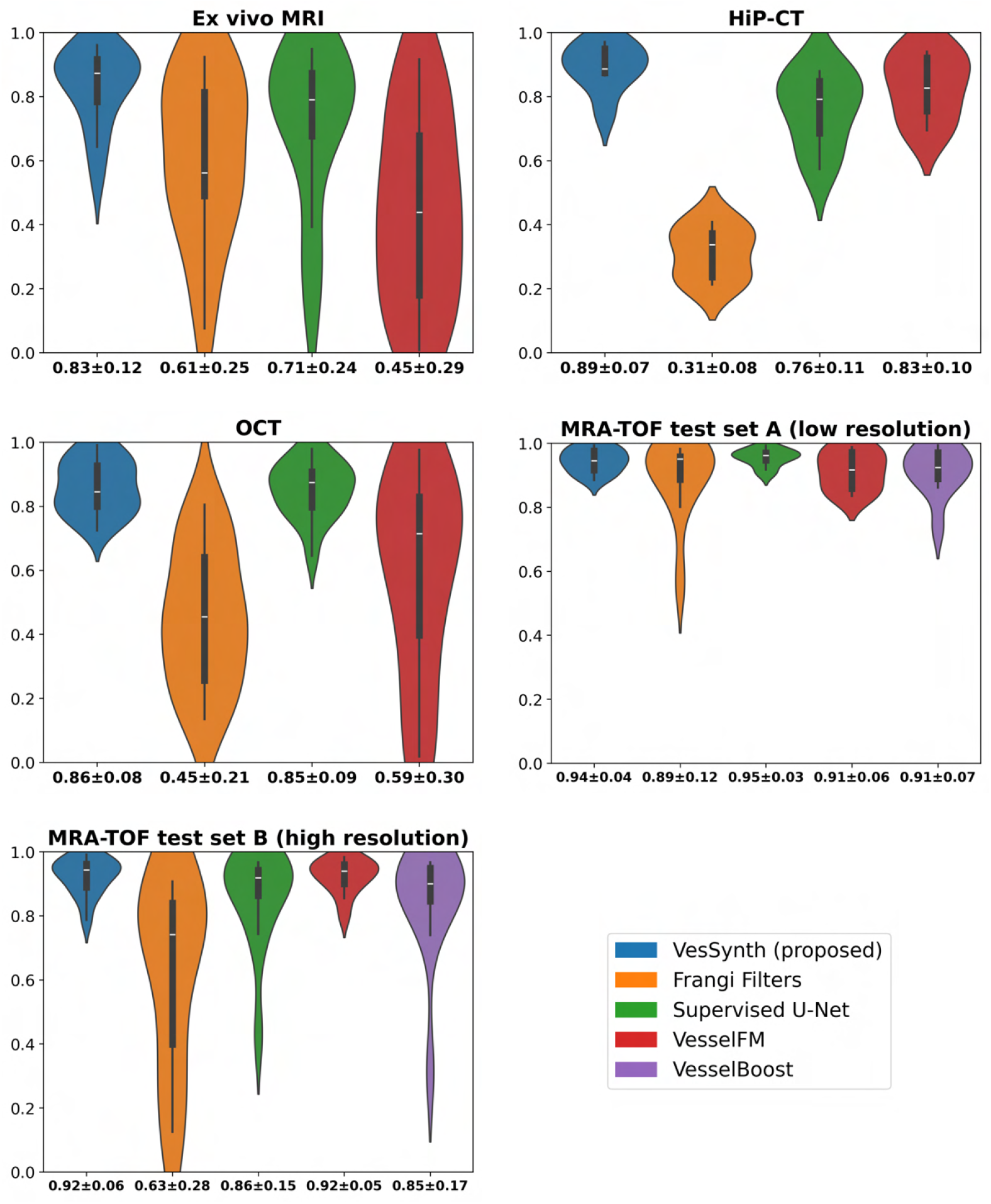
Violin plots of segmentation performance (corrected DSC) on real 64^3^-voxel test patches across modalities. VesSynth is compared with benchmark methods (Frangi filters, supervised U-Net, VesselFM, and VesselBoost for MRA-TOF). Each violin shows the distribution across patches, with an embedded boxplot (white median), and mean ± standard deviation reported below each violin plot.

Fig. 4 shows confusion maps for the proposed method and benchmarks on representative patches, illustrating true positives (green), false positives (red) and false negatives (yellow). MRA-TOF patches depict vascular structures spanning a range of calibers, with higher noise level in the high-resolution dataset; all methods detect both large and small vessels in these patches, although Frangi filters exhibit more errors. In *ex vivo*, VesselFM fails to detect both the vessel branching perpendicularly to the cortical surface and the vessel located within the white matter. Furthermore, the hypointense sulcus constitutes a challenging region and is erroneously segmented as a vessel by both Frangi filters and VesselFM. The HiP-CT patch depicts a large vessel with a clearly discernible lumen, vessel wall, and perivascular space, in addition to smaller vessels for which these structures are not resolved. Frangi filters, the supervised U-Net and VesselFM struggle to detect the smaller vessels, whereas the proposed method tends to slightly oversegment them. In OCT, a dark vessel is observed along with typical speckle noise artifacts. The proposed method correctly identifies the vessel but misses portions at the extremities. Its constraint toward hypointense vessels helps avoid false positives, whereas VesselFM, lacking this constraint, erroneously segments the bright shadow surrounding the vessel.

**Figure 4:**
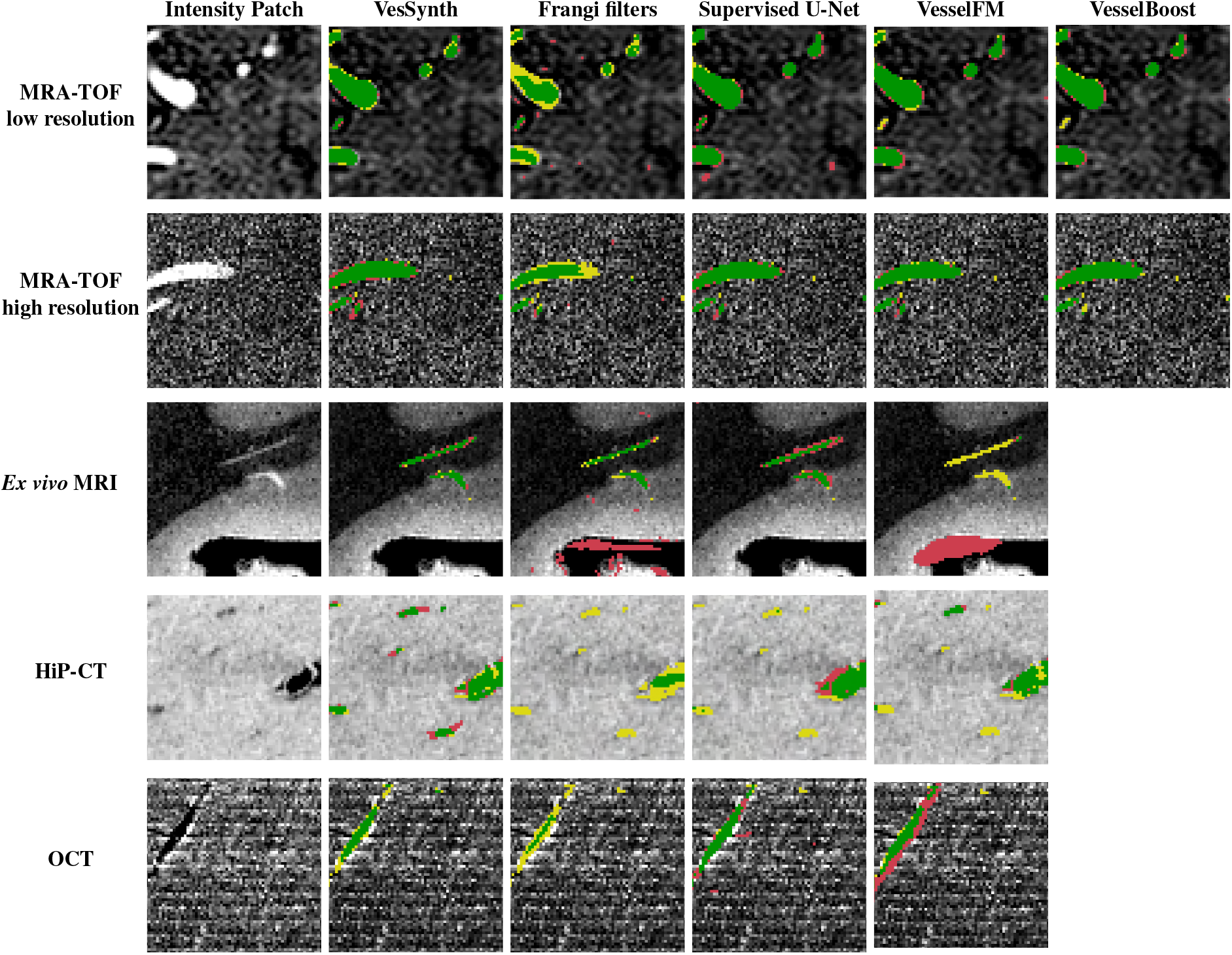
Confusion map of segmentations of proposed method and benchmarks, on a test patch for each modality. Green represents true positives, yellow represents false negatives and red denotes false positives.

### VesSynth scales to large unlabeled volumes

After benchmarking, we applied the proposed method to larger volumes lacking manual annotations, for qualitative assessment. Segmentations of a whole *ex vivo* MRI hemisphere, as well as whole OCT and HiP-CT sections are shown in Fig. 5. To facilitate visualization of the vascular architecture, projection of all vessels within a 100-voxel slab centered on the displayed intensity slice is shown, color-coded by distance from the slice. These visualizations highlight both the complexity of the segmentation task and the characteristic organization of the cerebral vasculature. The observed vascular architecture is consistent with known anatomical organization described in the literature [48]. In the cortex, vessels predominantly exhibit radial trajectories, with penetrating arterioles descending approximately perpendicular to the cortical surface and branching into a dense microvascular network, with typical near-orthogonal branch-ing patterns (blue circles in OCT image). Whole-hemisphere *ex vivo* MRI further reveals vessels in the white matter of the gyri coursing parallel to the gyral axes, extending toward and aligning along the outer boundary of ventricular walls, while deep medullary veins exhibit predominantly radial orientations [49]. The segmentations capture these known structural features, thereby supporting the biological plausibility of the proposed method.

**Figure 5:**
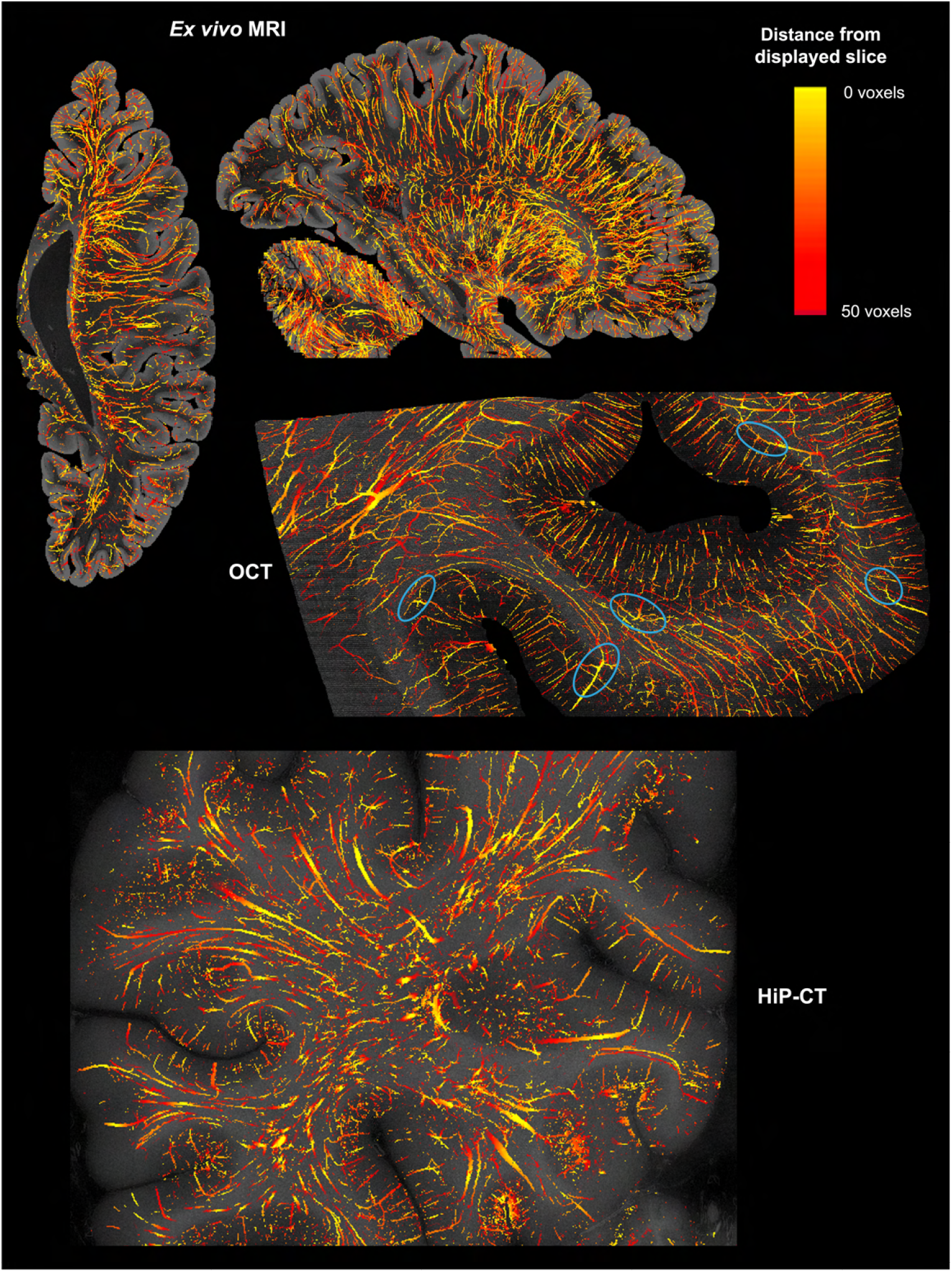
Vessel segmentation obtained by VesSynth on larger test volumes from *ex vivo* MRI (whole hemisphere), OCT (occipital cortex), and HiP-CT. The projection of all vessels contained in a 100-voxel thick slab centered around the displayed intensity slice is shown, with vessels color-coded by unsigned distance from the slice.

Fig. 6 shows vessel segmentation on a whole MRA-TOF image from test set A, for the proposed method and benchmarks. Given the reduced image size compared to the *ex vivo* modali-ties (attributable to the lower spatial resolution and slab-based acquisition), all vessels are projected and visualized in a single view, with color-coding indicating their distance from the central slice. Close-up visualizations demonstrate that both the proposed method and the supervised U-Net detect vessels across all calibers, whereas the other approaches fail to detect some of the smaller vessels. Overall, the proposed approach exhibits robust performance, consistently capturing vessels across multiple spatial scales.

**Figure 6:**
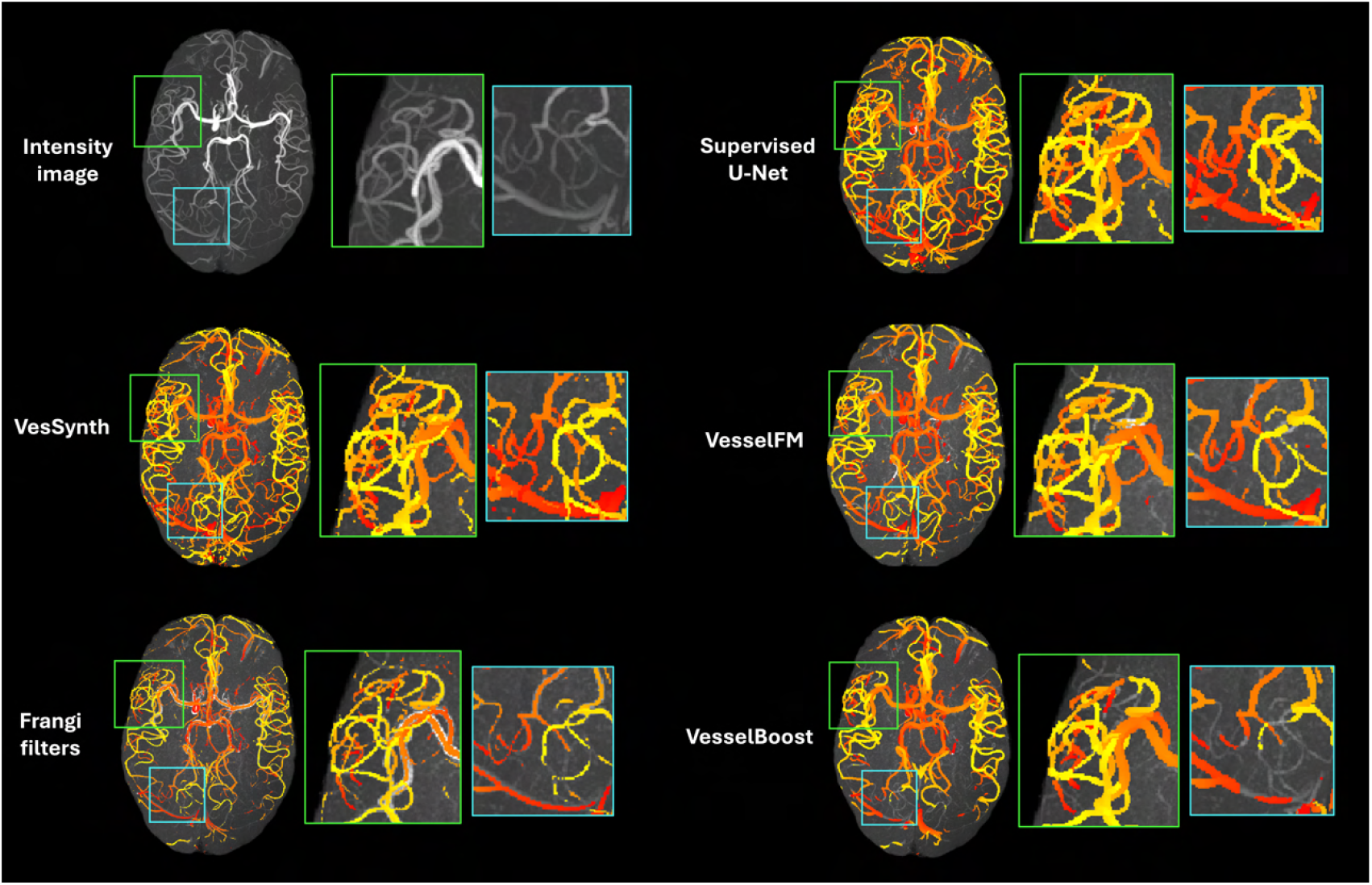
Vessel segmentations obtained by VesSynth and benchmark methods on a whole image from test set A. Maximum intensity projection of the original image is shown in the top left. The segmentations display all vessels within the image, color-coded by unsigned distance from the central slice. Enlarged views of two regions demonstrate that VesSynth effectively delineates vessels across a range of calibers, including the smallest.

## Discussion

In this work, we propose VesSynth, a framework for automatic vessel segmentation across multiple imaging modalities (*ex vivo* MRI, OCT, HiP-CT, and MRA-TOF) trained entirely on synthetic data. The proposed method achieves strong perfor-mance across all modalities (corrected DSC ranging from 0.83 to 0.94, with best performance in TOF-MRA), outperforming benchmark methods in most settings. Despite not being exposed to real data during training, the method enables accurate and robust segmentation on real images, highlighting the potential of synthetic-data–driven approaches. By splitting the TOF-MRA test set into subsets with different field strengths and resolutions, we further demonstrate the robustness of the proposed method to variations in imaging conditions within the same modality. Notably, VesselFM, which is trained on a mixture of real and synthetic data, performs well on modalities represented in its training set (HiP-CT and MRA-TOF) but less so on unseen modalities (OCT and *ex vivo* MRI), highlighting the importance of the proposed synthesis-based approach. For completeness, we note that VesselFM is applied out-of-the-box across modalities, whereas the proposed method and other benchmarks are optimized per modality.

As primary evaluation metric in this study we use corrected DSC, which discounts errors within a local neighborhood of true positives to account for the inherent ambiguity in vessel boundary delineation. The standard DSC sensitive to exact boundaries achieved by the proposed method ranges from 0.60 to 0.75 (Supplementary Table 3). The gap between standard and corrected DSC indicates that most discrepancies between automatic segmentation and manual labels arise from boundary uncertainty rather than missed detections. In terms of standard DSC, performance relative to benchmarks remains unchanged for *ex vivo* MRI and HiP-CT, where VesSynth continues to out-perform all other methods. On OCT and MRA-TOF, it drops by one rank: it ranks second to the supervised U-Net on OCT (0.60 vs 0.65); on MRA-TOF test set A, it ranks below the supervised U-Net (0.82) and VesselBoost (0.78), matching the performance of VesselFM (0.75); while on test set B, it ranks second behind VesselFM (0.78 vs 0.74).

A drawback of the proposed framework is the need to manually tune synthesis hyperparameters for each imaging modality, particularly vessel scale and contrast. Future work will focus on automating the synthesis to learn these parameters directly from data, incorporating topological constraints to preserve connectivity, and extending segmentation to vascular compartments and perivascular spaces. These developments are facilitated by the flexibility of the synthetic data paradigm, which enables tailored ground-truth generation.

VesSynth provides a flexible and scalable solution for vascular mapping across modalities and spatial scales. Because no single imaging modality captures the full range of vessel sizes, cross-modal segmentation is essential for comprehensive multiscale characterization of cerebrovascular architecture, which in humans has historically been achieved only through destructive vascular casting methods [48]. The proposed approach facilitates the construction of comprehensive vascular atlases, enabling the study of inter-individual variability and disease-related changes, ultimately supporting an exhaustive characterization of human cerebral angioarchitecture. This capability is particularly important given the growing recognition of vascular dysfunction as a central factor across brain disorders.

Beyond clinical relevance, vessel segmentation supports several methodological applications: (1) cross-modal image registration by using vessels as endogenous fiducial landmarks [50]; (2) slab stitching in serial-sectioning modalities through preservation of vessel continuity [51]; (3) white matter connectivity analysis by reducing misclassification of vessels as fiber pathways and enabling study of vessel–tract orientation relationships [52]; and (4) functional MRI analysis by distinguishing intra-vascular from extra-vascular contributions to the blood oxygenation level–dependent (BOLD) signal [53, 54, 55]. Finally, the synthetic training paradigm makes the framework extremely flexible, allowing easy extension to additional imaging modalities and histological techniques, as well as to other tubular structures such as axons and fiber bundles, whose mapping suffers from the same scaling limitations as cerebral vessels [56].

In conclusion, this work represents an important step toward scalable, cross-modal vascular analysis and provides a foundation for large-scale investigations of vascular contributions across a broad range of brain disorders, accelerating mechanistic and translational discoveries.

## Supporting information

This video shows 3D rendering of whole brain vessel segmentation

## Methods

### Synthetic vessels

Vascular trees were synthesized using a previously developed pipeline that generates vessels using a spline-based geometric model [39]. This synthetic framework imposes geometric, but not biophysical, constraints, providing maximal flexibility in structural configurations. Vascular tree generation is governed by the following parameters: voxel size; number of trees per unit volume (i.e., the expected number of vessels visible per unit volume in real images for that modality); number of offspring per parent spline; maximum tree depth (i.e. number of hierarchical levels in the tree); tortuosity (curvature degree); root vessel radius; branch radius factor (radius ratio between parent and child); and radius fluctuation along the spline. Parameter values are sampled from modality-specific distributions, as detailed in Table 1. For OCT, we adopted distributions optimized in prior work [39], whereas parameters were adapted for *ex vivo* MRI, HiP-CT and TOF-MRA to reflect the characteristics of each modality. Our statistical distributions intentionally extend beyond physiologically plausible ranges to expose the network to more diverse anatomical configurations during training than we expect at inference time in real data [40, 41].

**Table 1:** Distributions of synthesis parameters used to generate vascular trees across modalities. log 𝒩 (µ, σ) denotes a log-normal distribution, where µ and σ are mean and standard deviation of the corresponding Gaussian distribution. 𝒰 (*a, b*) is a continuous uniform distribution on the interval [*a, b*]. 𝒰_*z*_(*a, b*) is a discrete uniform distribution on the interval [*a, b*].

|  | *Ex vivo* MRI | HiP-CT | OCT | MRA-TOF |
| --- | --- | --- | --- | --- |
| Voxel size (isotropic) | 100μm | 30μm | 20μm | 400μm |
| Number of trees (per $mm^3$) | $\log\mathcal{N}(-4.952, 0.833)$ | $\mathcal{U}(0.03, 0.06)$ | $\mathcal{U}(0.1, 0.2)$ | $\mathcal{U}(1.25 * 10^{-5}, 2.5 * 10^{-5})$ |
| Number of offspring | $\log\mathcal{N}(1.263, 0.833)$ | $\mathcal{U}_z(1, 6)$ | $\mathcal{U}_z(1, 6)$ | $\mathcal{U}_z(1, 6)$ |
| Maximum tree depth | 2 | $\mathcal{U}_z(1, 6)$ | $\mathcal{U}_z(1, 6)$ | $\mathcal{U}_z(1, 6)$ |
| Tortuosity | $\log\mathcal{N}(1.456, 0.555)$ | $\mathcal{U}(1, 5)$ | $\mathcal{U}(1, 5)$ | $\mathcal{U}(1, 5)$ |
| Root radius (mm) | $\log\mathcal{N}(-2.322, 0.198)$ | $\log\mathcal{N}(-2.414, 0.472)$ | $\log\mathcal{N}(-2.322, 0.198)$ | $\log\mathcal{N}(0.682, 0.149)$ |
| Branch radius factor | $\log\mathcal{N}(-0.367, 0.142)$ | $\log\mathcal{N}(-0.296, 0.133)$ | $\log\mathcal{N}(-0.767, 0.385)$ | $\log\mathcal{N}(-0.442, 0.153)$ |
| Radius fluctuation factor | $\log\mathcal{N}(-0.005, 0.100)$ | $\log\mathcal{N}(-0.020, 0.198)$ | $\log\mathcal{N}(-0.020, 0.198)$ | $\log\mathcal{N}(-0.020, 0.198)$ |

**Table 2:** Summary of real datasets and extracted patches used for model validation and testing.

| Modality | Voxel size | # of different images | # of extracted patches | validation-test split<br># patches (# cases) |
| --- | --- | --- | --- | --- |
| *Ex vivo* MRI | 120-150 $\mu\text{m}$ isotropic | 12 | 37 | 22(7); 15(5) |
| HiP-CT | 25.25-30 $\mu\text{m}$ | 2 | 17 | 10(1); 7(1) |
| OCT | 20 $\mu\text{m}$ | 3 | 28 | 12(1); 16(2) |
| MRA-TOF | see Table 2b | 45 | 54 | 32(32); 22(13) |

| Dataset | Voxel size (mm) | # of extracted patches | Scanner | Use |
| --- | --- | --- | --- | --- |
| IXI-IOP | $0.26 \times 0.26 \times 0.80$ | 12 | GE 1.5T | Validation set |
| IXI-Guys | $0.47 \times 0.47 \times 0.80$ | 10 | Philips 1.5T | Validation set |
| IXI-HH | $0.47 \times 0.47 \times 0.80$ | 10 | Philips 3T | Test set A |
| TubeTK | $0.51 \times 0.51 \times 0.80$ | 10 | Siemens 3T | Validation set |
| Bollmann *et al.* [19] | 0.14–0.16 isotropic | 12 | Siemens 7T | Test set B |

(b) Summary of MRA-TOF datasets. The IXI dataset comprises images acquired at three sites (IOP, Guys, and HH) using different scanners. To account for differences in spatial resolution and field strength, the test set was split into test set A and test set B.

**Table 3:** Segmentation performance on real 64^3^-voxel test patches across modalities, comparing the proposed synthetic-trained model with benchmark methods (Frangi filters, supervised U-Net, VesselFM, and VesselBoost for MRA-TOF only). DSC = Dice score; AUC = area under the Receiver Operating Characteristic curve; TPR = true positive rate; FDR = false discovery rate.

| **Ex vivo MRI** |  |  |  |  |  |  |
| --- | --- | --- | --- | --- | --- | --- |
|  | Soft DSC | DSC | Corrected DCS | AUC | Corrected TPR | Corrected FDR |
| VesSynth (proposed) | 0.654 ± 0.139 | 0.610 ± 0.141 | 0.831 ± 0.117 | 0.961 ± 0.039 | 0.834 ± 0.156 | 0.158 ± 0.092 |
| Frangi filters | 0.336 ± 0.161 | 0.362 ± 0.180 | 0.610 ± 0.247 | 0.879 ± 0.046 | 0.768 ± 0.224 | 0.459 ± 0.283 |
| Supervised U-Net | 0.546 ± 0.201 | 0.508 ± 0.203 | 0.713 ± 0.242 | 0.923 ± 0.080 | 0.718 ± 0.202 | 0.268 ± 0.276 |
| VesselFM | - | 0.296 ± 0.213 | 0.445 ± 0.286 | - | 0.506 ± 0.289 | 0.484 ± 0.323 |

| **HiP-CT** |  |  |  |  |  |  |
| --- | --- | --- | --- | --- | --- | --- |
|  | Soft DSC | DSC | Corrected DSC | AUC | Corrected TPR | Corrected FDR |
| VesSynth (proposed) | 0.717 ± 0.097 | 0.659 ± 0.119 | 0.893 ± 0.069 | 0.980 ± 0.012 | 0.826 ± 0.118 | 0.015 ± 0.021 |
| Frangi filters | 0.272 ± 0.071 | 0.139 ± 0.053 | 0.311 ± 0.075 | 0.839 ± 0.066 | 0.186 ± 0.053 | 0.001 ± 0.002 |
| Supervised U-Net | 0.615 ± 0.098 | 0.572 ± 0.105 | 0.759 ± 0.109 | 0.914 ± 0.042 | 0.651 ± 0.111 | 0.073 ± 0.132 |
| VesselFM | - | 0.619 ± 0.148 | 0.830 ± 0.095 | - | 0.723 ± 0.138 | 0.003 ± 0.002 |

| **OCT** |  |  |  |  |  |  |
| --- | --- | --- | --- | --- | --- | --- |
|  | Soft DSC | DSC | Corrected DSC | AUC | Corrected TPR | Corrected FDR |
| VesSynth (proposed) | 0.605 ± 0.160 | 0.596 ± 0.157 | 0.859 ± 0.081 | 0.978 ± 0.021 | 0.875 ± 0.113 | 0.132 ± 0.127 |
| Frangi filters | 0.316 ± 0.046 | 0.169 ± 0.103 | 0.451 ± 0.209 | 0.885 ± 0.060 | 0.316 ± 0.185 | 0.003 ± 0.011 |
| Supervised U-Net | 0.674 ± 0.103 | 0.646 ± 0.100 | 0.851 ± 0.085 | 0.986 ± 0.010 | 0.908 ± 0.059 | 0.191 ± 0.123 |
| VesselFM | - | 0.402 ± 0.261 | 0.587 ± 0.305 | - | 0.599 ± 0.247 | 0.220 ± 0.310 |

| **MRA-TOF** |  |  |  |  |  |  |
| --- | --- | --- | --- | --- | --- | --- |
|  | Soft DSC | DSC | Corrected DSC | AUC | Corrected TPR | Corrected FDR |
| VesSynth (proposed) - test set A | 0.791 ± 0.084 | 0.750 ± 0.096 | 0.945 ± 0.035 | 0.993 ± 0.004 | 0.919 ± 0.060 | 0.024 ± 0.038 |
| VesSynth (proposed) - test set B | 0.768 ± 0.094 | 0.741 ± 0.104 | 0.922 ± 0.056 | 0.982 ± 0.014 | 0.901 ± 0.076 | 0.050 ± 0.066 |
| Frangi filters - test set A | 0.533 ± 0.110 | 0.555 ± 0.094 | 0.893 ± 0.120 | 0.912 ± 0.022 | 0.935 ± 0.035 | 0.122 ± 0.178 |
| Frangi filters - test set B | 0.321 ± 0.135 | 0.321 ± 0.167 | 0.628 ± 0.278 | 0.885 ± 0.043 | 0.798 ± 0.117 | 0.341 ± 0.367 |
| Supervised U-Net - test set A | 0.850 ± 0.049 | 0.825 ± 0.052 | 0.955 ± 0.026 | 0.993 ± 0.004 | 0.926 ± 0.043 | 0.014 ± 0.015 |
| Supervised U-Net - test set B | 0.673 ± 0.154 | 0.627 ± 0.155 | 0.859 ± 0.147 | 0.974 ± 0.049 | 0.850 ± 0.181 | 0.094 ± 0.108 |
| VesselFM - test set A | - | 0.751 ± 0.120 | 0.914 ± 0.057 | - | 0.852 ± 0.094 | 0.009 ± 0.009 |
| VesselFM - test set B | - | 0.777 ± 0.078 | 0.924 ± 0.052 | - | 0.883 ± 0.077 | 0.027 ± 0.047 |
| VesselBoost - test set A | - | 0.781 ± 0.129 | 0.914 ± 0.073 | - | 0.852 ± 0.117 | 0.004 ± 0.004 |
| VesselBoost - test set B | - | 0.676 ± 0.174 | 0.848 ± 0.173 | - | 0.768 ± 0.198 | 0.018 ± 0.049 |

After sampling parameters from these distributions, the resulting vascular trees were rasterized to produce two outputs: (1) a label volume ***l*** in which each vessel is assigned a unique identifier to preserve vessel-level correspondence during synthetic image generation, and (2) a probability volume ***p*** encoding the likelihood of each voxel belonging to the vasculature to model partial volume effects. The latter represents an extension of previous work [39], in which probabilistic information was not utilized. Full details about the rasterization procedure are provided in [39].

In HiP-CT, distinct vascular compartments (lumen, wall, and perivascular space) are visible, which required the synthesis framework to be extended. Vessels were eroded twice using smooth spatial fields, once to generate a perivascular layer and once to form the vessel wall. Then, separate identifiers were assigned to the resulting compartments, while the probability volume remained unchanged.

For each modality, 1,000 synthetic images with 128 ×128 ×128 voxels were generated, of which 800 were used for training and 200 for validation on synthetic data.

### Synthetic images

We developed a modality-specific pipeline to generate synthetic intensity images corresponding to given vascular trees. The parenchyma was modeled using random smooth spline-based shapes as in [44, 39], while a vascular volume was initialized using the vascular trees in the label map ***l***. Random Gaussian-distributed intensities were then assigned to vessels and parenchymal structures, and the two volumes were fused into a composite image, which subsequently underwent a sequence of intensity augmentations. Specific details of the synthesis procedure varied across imaging modalities, aiming to capture modality-dependent features, such as vessel-to-parenchyma contrast, noise characteristics, and acquisition ar-tifacts, while intentionally extending beyond realistic ranges to promote generalizability. A detailed description of the modality-specific intensity synthesis is provided below and examples are shown in Fig. 2. In the remainder, ***y*** denotes the parenchymal image, ***𝓏*** the the vascular image, and ***x*** the composite image. 𝒩 (µ, σ) denotes a Gaussian distribution with mean µ and standard deviation σ; 𝒰 (*a, b*) denotes a continuous uniform distribution on [*a, b*]; 𝒰_*z*_(*a, b*) is a discrete uniform distribution over integers in [*a, b*]; and ∼ indicates sampling.

#### *Ex vivo* MRI

provides non-specific contrast for vessels, which may appear brighter or darker than surrounding tissue, and the same vessel may change intensity along its course. The physical origins of this signal variability remain uncertain, with potential contributors including residual blood, iron deposition, fixation artifacts, or microstructural voids. To simulate this phenomenon, after initializing the vessel intensity volume ***𝓏*** using the synthetic vessels in ***l***, we injected in ***𝓏*** smooth random structures, which were then randomly grouped into *N* classes, with *N*∼𝓊 _*z*_(1, 5). All shapes in class *i* (*i* = 1, …, *N*) were assigned a shared ID and intersected with the vessel trees, which inherited the the same IDs as the intersecting shapes, while regions outside the vasculature were discarded. This enabled individual vessels to smoothly inherit multiple IDs along their trajectory. Vessel intensities were then sampled using a Gaussian Mixture Model (GMM): for each vessel class *i*, intensities were drawn from 𝒩 (µ_*i*_, σ_*i*_) with µ_*i*_∼𝒰 (0, 1), σ_*i*_∼ *𝒰* (0, 0.05), *i* = 1, …, *N*.

The volume ***y*** with parenchymal tissue was synthesized by generating *M* smooth random shapes with *M*∼ *𝒰*_*z*_(1, 16), and random spherical structures with radius ∼𝒰_*z*_(0, 5) voxels and frequency ∼𝒰 (0, 0.03). Intensities were then drawn for each parenchymal structure *j* using a Gaussian Mixture Model with 𝒩 (µ _*j*_, σ _*j*_) with µ _*j*_ ∼ 𝒰 (0, 1), σ _*j*_ ∼ 𝒰 (0, 0.05) ∀ *j*.

The foreground volume ***𝓏*** and the background volume ***y*** were then merged: ***𝓏*** was rescaled to [0, 1], ***y*** was rescaled to [*a*, 1] with *a*∼*𝒰* (0.2, 0.4), and the two were combined into a final intensity volume ***x*** = ***p𝓏***+(1− ***p***)***y***, where ***p*** denotes the vessel probability field. In ***x***, vessels can appear brighter, darker, or indistinguishable from the background, with no contrast constraints imposed, and intensity can vary along a vessel centerline. Finally, a set of intensity augmentations was applied to ***x***: gamma transform with exponent ∼𝒰 (0, 5); a random multiplicative smooth bias field with values in (0, 1); Gaussian smoothing with full-width at half-maximum ∼𝒰 (0, 2); and random non-central Chi noise with standard deviation ∼ 𝒰 (0, 0.03).

#### HiP-CT

displays hypointense vessels, with larger vessels often exhibiting compartmental structure with a dark lumen, a brighter wall, and a darker perivascular space. We sought to generate vessels darker than the surrounding parenchymal tissue, but only relative to their immediate neighborhood rather than globally across the volume. To this end, we defined the vessel intensity volume ***𝓏*** as a voxel-wise transfor-mation of the parenchymal intensity volume ***y***:

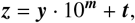

where ***m*** and ***t*** are synthesized modulation fields, and multiplication and exponentiation are applied element-wise. By restricting the allowable ranges of ***m*** and ***t***, we impose the desired contrast between ***𝓏*** and ***y*** at corresponding voxels; because ***y*** varies smoothly, this constraint extends to the surrounding neighborhood in ***y***.

The background volume ***y*** was synthesized following the same procedure as for *ex vivo* MRI, and then rescaled to [*a*, 1] with *a*∼ *𝒰* (0.2, 0.4). The modulation fields ***m*** and ***t*** were initialized using the synthetic vessels in the label volume ***l***, with each vessel (and compartment, when visible) carrying a unique identifier. All structures in ***m*** were randomly grouped into *N*_1_ classes, with *N*_1_ ∼ 𝒰_*z*_(1, 8), and class-specific intensities were sampled from 𝒩 (µ_*i*_, σ_*i*_), where µ_*i*_ ∼ *𝒰* (0, 1) and σ_*i*_ ∼ *𝒰* (0, 0.05), for *i* = 1, …, *N*_1_. Because grouping was random, distinct compartments of the same vessel could be assigned to the same or to different classes. The field ***m*** was then rescaled to [*b, c*], with *b* ∼ 𝒰 (0.08, 0.24), *c* ∼ 𝒰 (0.3, 0.6), and exponentiated voxel-wise to yield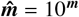. This ensures that the modulation factor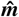 applied to ***y*** varies, at most, in the range [1.2, 4] approximately. Similarly, all structures in ***t*** were randomly grouped into *N*_2_ classes, with *N*_2_ ∼𝒰_*z*_(1, 8), and class-specific intensities were sampled from a Gaussian mixture (µ_*i*_, σ_*i*_), where µ_*i*_ ∼ *𝒰* (0, 1) and σ_*i*_∼ *𝒰* (0, 0.01), for *i* = 1, …, *N*_2_. The field ***t*** was subsequently rescaled to [0, 1].

We then computed ***𝓏*** = ***y***· 10^***m***^ + ***t*** and fused it with the parenchymal background ***y*** to obtain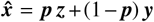. With the chosen parameter ranges, vessels in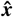 exhibited higher intensities than their immediate surroundings. To model the dark vessel appearance seen in HiP-CT, we inverted the contrast by computing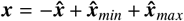. With this synthesis procedure, vessels in ***x*** are hypointense relative to their local parenchymal surroundings, and compartments of the same vessel may share or differ in intensity classes. When intensities differ, no constraint is imposed on their relative appearance, except that all compartments remain darker than their local background. This behavior does not reproduce realistic compartmental contrast patterns, but the synthesis is intentionally extended beyond realism to enhance generalization.

As final set of intensity augmentations, we applied: gamma transform with exponent ∼𝒰 (0, 5); a random multiplicative smooth bias field with values in (0, 1); Gaussian smoothing with full-width at half-maximum ∼𝒰 (0, 2); random noise, applied with equal probability as either non-central Chi noise (standard deviation ∼ 𝒰 (0, 0.15)) or multiplicative Gamma noise (mean=1, standard deviation ∼ 𝒰 (0, 0.1)).

#### OCT

exhibits hypointense vessels due to their low coherent backscatter relative to surrounding tissue. We adopted a synthesis pipeline similar to HiP-CT, with vessels darker than the surrounding parenchyma in their immediate neighborhood. The background volume ***y*** was synthesized and rescaled as for HiP-CT, but with increased variability in random shapes (*M* ∼*𝒰*_*z*_(1, 18)). Modulation fields ***m*** and ***t*** also followed the same framework as HiP-CT, with adjusted parameters. To generate ***m***, vessels were grouped into *N*_1_ ∼ *𝒰*_*z*_(1, 5) classes and class-specific intensities were sampled from 𝒩 (µ_*i*_, σ_*i*_), where µ_*i*_ ∼ *𝒰* (0, 1) and σ_*i*_ ∼ *𝒰* (0, 0.05). The field ***m*** was then rescaled to [*b, c*], with *b* ∼ *𝒰* (0.1, 0.3) and *c* ∼ *𝒰* (0.4, 0.8), and exponentiated to yield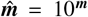, corresponding to modulation factors of approximately [1.25, 6.3]. Similarly, ***t*** was constructed by grouping vessels into *N*_2_∼ 𝒰_*z*_(1, 5) classes with intensities sampled from 𝒩 (µ_*i*_, σ_*i*_) with µ_*i*_∼𝒰 (0, 1), σ_*i*_∼𝒰 (0, 0.01), and then rescaled to [0.3, 1].

The combined intensity volume ***x*** was computed as in HiP-CT. Augmentations also followed the same procedure, with two modifications: (1) a slab-wise multiplicative bias field with slab thickness ∼𝒰_*z*_(1, 4) was applied before smoothing to simulate slab-wise *z*-decay in OCT; and (2) to model speckle noise, we applied to each patch a multiplicative Gamma noise with mean ∼ 𝒰 (0.5, 1) and standard deviation∼ 𝒰 (0, 0.5).

#### MRA-TOF

After initializing the vessel intensity volume ***𝓏*** using the synthetic vessels in ***l***, each containing a unique ID, vessels were randomly grouped into *N* classes with *N* ∼𝒰_*z*_(1, 5), and class-specific intensities were sampled from 𝒩 (µ_*i*_, σ_*i*_) with µ_*i*_ ∼*𝒰* (0, 1), σ_*i*_ ∼*𝒰* (0, 0.05), for each class *i* = 1, …, *N*.

The parenchymal volume ***y*** was synthesized by generating *M* smooth random shapes, with *M*∼*𝒰* _*z*_(1, 18), and random spherical structures whose radii were drawn from *𝒰*_*z*_(0, 5) voxels and occurrence frequency from (0, 0.03). Intensities were then drawn for each parenchymal structure *j* using a Gaussian Mixture Model with 𝒩 (µ _*j*_, σ _*j*_) with µ _*j*_ ∼*𝒰* (0, 1), σ _*j*_ ∼ *𝒰* (0, 0.05) ∀ *j*.

To simulate inflow-related bright vessel appearance characteristic of MRA-TOF imaging, ***𝓏*** was rescaled to [*b, c*], with *b*∼ *𝒰* (1, 1.5) and *c* ∼ *𝒰* (2, 3), while ***y*** was rescaled to [*a*, 1], with *a* ∼ *𝒰* (0.2, 0.4), before fusing them into a final intensity image ***x*** = ***p𝓏*** + (1 −***p***)***y***. This ensures that vessels remained brighter than surrounding tissue. The resulting volume subsequently underwent a set of augmentations: Perlin-like noise with intensities ~ 𝒰 (0.3, 1.5) applied to each patch with probability 70%; random spherical structures with radius ∼ 𝒰 (2, 5) voxels and intensities ~ 𝒰 (1, 1.8); gamma transform with exponent ∼𝒰 (0, 5); a random multiplicative smooth bias field with values in (0, 1); Gaussian smoothing with full-width at half-maximum ∼ 𝒰 (0, 2); and random non-central Chi noise with standard deviation ∼ 𝒰 (0, 0.06).

For all modalities, synthetic intensity volumes were normalized at the end of the synthesis pipeline by mapping the 1^*st*^ and 99^*th*^ percentiles to 0 and 1, respectively.

### Real data and extraction of patches

We collected real images from all modalities presented in this paper, from which small volumetric patches (64^3^ voxels) were extracted and manually labeled. The small size of these validation patches ensures exhaustive annotation and avoids being biased by the operators’ fatigue. This resulted in the following dataset:

#### Ex vivo MRI

12 *ex vivo* MRI hemispheres were obtained from the Massachusetts General Hospital Autopsy Suite and scanned *in situ* using a multi-echo fast low-angle shot sequence. Image resolution ranged from 120 µm to 150 µm, with acquisitions performed on either 3T or 7T Siemens scanners. Full protocol details are reported in [50], and a subset of hemispheres is publicly available on the DANDI platform (DANDI:000026) at https://dandiarchive.org/dandiset/000026. Two to five patches were extracted from each hemisphere across distinct anatomical regions, yielding a total of 37 patches.

#### HiP-CT

We used two publicly available HiP-CT scans: LADAF-2021-17, available on the Human Organ Atlas [57] (http://doi.org/10.15151/ESRF-DC-1773964937) and I58, available on the DANDI platform (DANDI:000026) at https://dandiarchive.org/dandiset/000026. LADAF-2021-17 has a native resolution of 25.25 µm and 10 patches were extracted. I58 has a native isotropic resolution of 15 µm, which was downsampled to 30 µm before extracting 7 patches.

#### OCT

We used the datasets described in [39], obtained from the Massachusetts General Hospital Autopsy Suite and imaged locally. These include a sample of primary somatosensory cortex imaged by a 1300 nm spectral domain serial-section OCT system, with an in-plane resolution of 10 µm and an axial resolution of 3.5 µm [58] (sample I46, available on the DANDI platform at https://dandiarchive.org/dandiset/000722), and both frontal and occipital cortices of two other locally-acquired samples imaged by a different 1300 nm polarization sensitive serial-section OCT system [59]. All volumes were reconstructed to an isotropic resolution of 20 µm [60]. We extracted 12 random 64^3^ patches from the somatosensory dataset I46, and 8 patches from each of the other two datasets (four from the frontal cortex and four from the occipital cortex per case), for a total of 28 patches.

#### MRA-TOF

We compiled MRA-TOF images from multiple datasets: the IXI dataset (http://brain-development.org/ixi-dataset/), which includes acquisitions from three sites using different scanners; the TubeTK dataset (https://data.kitware.com/api/v1/folder/58a3abaa8d777f0721a65b0a/download); and a set of high-resolution scans (https://doi.org/10.17605/OSF.IO/NR6GC) [19]. A random subset of scans were selected from the IXI (32 scans) and TubeTK (10 scans), and one 64^3^ patch was extracted from each image. Three images from the high-resolution dataset were used, with four patches extracted from each images. In total, 45 images were included and 54 patches were extracted. Details on image resolution and scanner characteristics are summarized in Table 2b.

For each modality, we divided the extracted patches into a validation set, used for model selection, and a test set, used for independent evaluation. To avoid information leaks, when multiple patches had been extracted from the same volume, all were assigned to the same split (either validation or test). For MRA-TOF, the test set was further subdivided into test set A and test set B to account for differences in spatial resolution and field strength (Table 2b). A summary of all extracted patches and dataset splits is provided in Table 2.

### Vessel manual labeling

Vessels were manually labeled in all patches. Given the difficulty of exhaustively annotating vascular structures, we established a structured labeling protocol involving two scenarios:

#### Expert raters

Experienced raters annotated the patches sequentially to ensure completeness. Typically, two passes were performed by two different raters; for the most challenging patches, a third pass was conducted by an additional expert.

#### Junior raters under expert supervision

Some patches were annotated by raters without prior experience in vessel labeling, under close expert supervision and guidance. They junior raters first completed a training phase, during which they annotated a test volume and compared their results with expert annotations. Only raters achieving a Dice score above 0.6 were allowed to proceed to the next phase. Each patch was then independently annotated by two raters, with periodic feedback provided by an expert. A final label was defined as the union of the two annotations, as omissions were more frequent than false positives, and was subsequently reviewed by an expert to ensure completeness.

All labeling was performed using FreeView [61] with the voxel-edit tool (brush size 1–3 voxels). Raters adjusted image contrast to optimize vessel visibility and used multiple orthogonal views to identify tubular structures. Examples of manual vessel annotations are shown in Fig.1.

### Neural network and model training

For each modality, we trained a neural network on synthetic data using a 3D U-Net architecture [46] with residual connections, comprising five resolution levels and two 3× 3× 3 convolutions per level. The model was trained using a soft Dice loss, a learning rate of 10^−2^, and a batch size of one. Training was performed for 500 epochs, with each epoch encompassing all 800 pre-computed synthetic vascular trees.

At each training step, a synthetic label volume ***l***, assigning a unique identifier to each vascular tree, and the corresponding probability map ***p*** were randomly sampled and augmented through flipping, elastic deformation, and smooth spatial masking. An intensity image was then synthesized on-the-fly using the modality-specific pipeline, avoiding repeated exposure to identical examples, as opposed to vessel label synthesis and rasterization, which was pre-computed due to its higher computational cost. The resulting intensity volume was fed to the network for training. Although both ***l*** and ***p*** were used during intensity synthesis, only ***p*** served as ground truth in the loss computation.

Model performance was monitored across epochs using Dice scores on both the synthetic validation set (200 samples) and the real validation set of labeled 64^3^-voxel patches. We ensured that the training reached convergence on both validation sets and the final model was selected as the epoch achieving the highest Dice score on the real validation set.

### Benchmarking and validation metrics

We compared the proposed method to the following benchmarks, tested on the same test set as the proposed method:

#### Supervised U-Nets trained on real data

For each modality, we trained a supervised U-Net on real annotated data to assess whether it provides an advantage over the use of synthetic images. The supervised networks used the same architecture as the proposed method. Here, the validation set used previously was further split into separate training and validation sets. For each modality, the resulting splits were as follows: *ex vivo* MRI, 11 training and 11 validation patches; HiP-CT, 5 training and 5 validation patches; OCT, 7 training and 5 validation patches; and MRA-TOF, 22 training and 10 validation patches. Validation sets were used to monitor performance across epochs and to select the model achieving the highest Dice score. Given the limited amount of training data, extensive data augmentation was applied, including geometric transformations (random flipping and elastic deformation) and intensity augmentations (gamma transformation, multiplicative bias fields, Gaussian smoothing, and Gaussian noise)

#### Frangi filters

a modality-agnostic method for detecting tubular structures based on the eigenvalues of the Hessian matrix [24]. Hyperparameters, including sensitivity to platelike and blob-like structures and the scale, were tuned separately for each modality using the same validation set as for the proposed method.

#### VesselFM

a Deep Learning foundation model recently developed for vessel segmentation [38], trained on a mixture of real and synthetic data spanning multiple modalities, organs, and species. The real training data include MRA-TOF, HiP-CT, computed tomography, X-ray, light-sheet microscopy, and multiphoton microscopy. The model is modality-agnostic, requires no hyperparameter tuning, and was directly evaluated on the test set.

#### VesselBoost

a Deep Learning method developed for MRA-TOF vessel segmentation [32]. We compared it with the proposed method on MRA-TOF data. No hyperparameter tuning was required, and the model was directly evaluated on the test set, after bias-field correction and non-local means denoising [62], as required by its pipeline Segmentation accuracy was evaluated based on the following metrics: *Dice score* (DSC) [47], computed from binarized predictions obtained by thresholding the probability map at 0.5; *soft* DSC, computed from continuous predictions in [0, 1]; *corrected* DSC; area under the receiver operating characteristic curve (AUC); *corrected* true positive rate (TPR); and *corrected* false discovery rate (FDR). We defined the corrected DSC as an extension of the standard DSC that disregards errors within a local neighborhood of true positives, defined as voxels sharing a face up to two adjacency steps. This metric de-emphasizes small boundary discrepancies near true positives, thereby assessing vessel localization while reducing sensitivity to exact boundary placement. This choice is motivated by the observation that vessel boundaries can be subjective even among expert raters. Similarly, corrected TPR and FDR are computed by ignoring errors within the same local neighborhood.

## Acknowledgements

Much of the computational resources required for this research was generously provided by the Massachusetts Life Sciences Center. We thank Patrick R. Hof for valuable input on cerebrovascular anatomy.

## Code availability

Code available at https://github.com/chiara-mauri/VesSynth.git

## Data availability

A subset of *ex vivo* MRI hemispheres is publicly available on the DANDI platform (DANDI:000026) at https://dandiarchive.org/dandiset/000026. HiP-CT scans are publicly available: LADAF-2021-17 is available on the Human Organ Atlas [57] (http://doi.org/10.15151/ESRF-DC-1773964937) and I58 is available on the DANDI platform (DANDI:000026) at https://dandiarchive.org/dandiset/000026. The OCT dataset I46 is available on the DANDI platform (DANDI:000722) at https://dandiarchive.org/dandiset/000722. The MRA-TOF images are publicly available: the IXI dataset at http://brain-development.org/ixi-dataset/; the TubeTK dataset at https://data.kitware.com/api/v1/folder/58a3abaa8d777f0721a65b0a/download); the high-resolution scans from [19] are available at https://doi.org/10.17605/OSF.IO/NR6GC.

## References

[1] Li, Q. et al. Cerebral small vessel disease. Cell transplantation 27, 1711–1722 (2018).

[2] Noto, N. M., Speth, R. C. & Robison, L. S. Cerebral amyloid angiopathy: a narrative review. Frontiers in Aging Neuroscience 17, 1632252 (2025).

[3] Mestre, H., Kostrikov, S., Mehta, R. I. & Nedergaard, M. Perivascular spaces, glymphatic dysfunction, and small vessel disease. Clinical science 131, 2257–2274 (2017).

[4] Yu, L., Hu, X., Li, H. & Zhao, Y. Perivascular spaces, glymphatic system and mr. Frontiers in Neurology 13, 844938 (2022).

[5] Debette, S., Schilling, S., Duperron, M.-G., Larsson, S. C. & Markus, H. S. Clinical significance of magnetic resonance imaging markers of vascular brain injury: a systematic review and meta-analysis. JAMA neurology 76, 81–94 (2019).

[6] Wardlaw, J. M., Smith, C. & Dichgans, M. Small vessel disease: mechanisms and clinical implications. The Lancet Neurology 18, 684–696 (2019).

[7] Bos, D. et al. Cerebral small vessel disease and the risk of dementia: a systematic review and meta-analysis of population-based evidence. Alzheimer’s & Dementia 14, 1482–1492 (2018).

[8] Fang, Y. et al. Cerebral small-vessel disease and risk of incidence of depression: A meta-analysis of longitudinal cohort studies. Journal of the American Heart Association 9, e016512 (2020).

[9] Qiu, W. Q. et al. Effects of white matter integrity and brain volumes on late life depression in the framingham heart study. International journal of geriatric psychiatry 32, 214–221 (2017).

[10] Chen, Y. et al. Correlation between cerebral small vessel disease and late-life depression: Insights from neuroimaging studies. Brain Research 1868, 150001 (2025).

[11] Korten, N. C., Comijs, H. C., Lamers, F. & Penninx, B. W. Early and late onset depression in young and middle aged adults: differential symptomatology, characteristics and risk factors? Journal of affective disorders 138, 259–267 (2012).

[12] Hoglund, Z. et al. Brain vasculature accumulates tau and is spatially related to tau tangle pathology in alzheimer’s disease. Acta Neuropathologica 147, 101 (2024).

[13] Rodríguez, J. L. et al. Entorhinal vessel density correlates with phosphorylated tau and tdp-43 pathology. Alzheimer’s & Dementia 20, 4649 (2024).

[14] Sweeney, M. D., Kisler, K., Montagne, A., Toga, A. W. & Zlokovic, B. V. The role of brain vasculature in neurodegenerative disorders. Nature neuroscience 21, 1318–1331 (2018).

[15] Carmeliet, P. Angiogenesis in health and disease. Nature medicine 9, 653–660 (2003).

[16] Carmeliet, P. & Jain, R. K. Molecular mechanisms and clinical applications of angiogenesis. Nature 473, 298–307 (2011).

[17] Zlokovic, B. V. The blood-brain barrier in health and chronic neurode-generative disorders. Neuron 57, 178–201 (2008).

[18] Todorov, M. I. et al. Machine learning analysis of whole mouse brain vasculature. Nature methods 17, 442–449 (2020).

[19] Bollmann, S. et al. Imaging of the pial arterial vasculature of the human brain in vivo using high-resolution 7t time-of-flight angiography. Elife 11, e71186 (2022).

[20] Gulban, O. F. et al. Whole-brain meso-vein imaging in living humans using fast 7-t mri. Science advances 12, eaea4540 (2026).

[21] Edlow, B. L. et al. 7 tesla mri of the ex vivo human brain at 100 micron resolution. Scientific data 6, 244 (2019).

[22] Cassot, F., Lauwers, F., Fouard, C., Prohaska, S. & Lauwers-Cances, V. A novel three-dimensional computer-assisted method for a quantitative study of microvascular networks of the human cerebral cortex. Microcirculation 13, 1–18 (2006).

[23] Reina-De La Torre, F., Rodriguez-Baeza, A. & Sahuquillo-Barris, J. Morphological characteristics and distribution pattern of the arterial vessels in human cerebral cortex: a scanning electron microscope study. The Anatomical Record: An Official Publication of the American Association of Anatomists 251, 87–96 (1998).

[24] Frangi, A. F., Niessen, W. J., Vincken, K. L. & Viergever, M. A. Multiscale vessel enhancement filtering. In International conference on medical image computing and computer-assisted intervention, 130–137 (Springer, 1998).

[25] Zana, F. & Klein, J.-C. Segmentation of vessel-like patterns using mathematical morphology and curvature evaluation. IEEE transactions on image processing 10, 1010–1019 (2001).

[26] Dokládal, P., Lohou, C., Perroton, L. & Bertrand, G. Liver blood vessels extraction by a 3-d topological approach. In International Conference on Medical Image Computing and Computer-Assisted Intervention, 98–105 (Springer, 1999).

[27] Yang, J. et al. Volumetric characterization of microvasculature in ex vivo human brain samples by serial sectioning optical coherence tomography. IEEE Transactions on Biomedical Engineering 69, 3645–3656 (2022).

[28] Yousefi, S., Liu, T. & Wang, R. K. Segmentation and quantification of blood vessels for oct-based micro-angiograms using hybrid shape/intensity compounding. Microvascular research 97, 37–46 (2015).

[29] Longo, A. et al. Assessment of hessian-based frangi vesselness filter in optoacoustic imaging. Photoacoustics 20, 100200 (2020).

[30] Goni, M. R., Ruhaiyem, N. I. R., Mustapha, M., Achuthan, A. & Nassir, C. M. N. C. M. Brain vessel segmentation using deep learning—a review. IEEE access 10, 111322–111336 (2022).

[31] Moccia, S., De Momi, E., El Hadji, S. & Mattos, L. S. Blood vessel segmentation algorithms—review of methods, datasets and evaluation metrics. Computer methods and programs in biomedicine 158, 71–91 (2018).

[32] Xu, M. et al. Vesselboost: A Python Toolbox for Small Blood Vessel Segmentation in Human Magnetic Resonance Angiography Data. Aperture Neuro 4 (2024).

[33] Yagis, E. et al. Deep learning for 3d vascular segmentation in hierarchical phase contrast tomography: a case study on kidney. Scientific Reports 14, 27258 (2024).

[34] Kaplan, J. et al. Scaling laws for neural language models. arXiv preprint 2001.08361 (2020).

[35] Kirillov, A. et al. Segment anything. In Proceedings of the IEEE/CVF international conference on computer vision, 4015–4026 (2023).

[36] Ma, J. et al. Segment anything in medical images. Nature communications 15, 654 (2024).

[37] Holroyd, N. A. et al. tubenet: a generalizable deep learning tool for 3d vessel segmentation. Biology Methods and Protocols 10, bpaf087 (2025).

[38] Wittmann, B., Wattenberg, Y., Amiranashvili, T., Shit, S. & Menze, B. vesselfm: A foundation model for universal 3d blood vessel segmentation. In Proceedings of the Computer Vision and Pattern Recognition Conference, 20874–20884 (2025).

[39] Chollet, E. et al. Neurovascular segmentation in soct with deep learning and synthetic training data. arXiv preprint 2407.01419 (2024).

[40] Gopinath, K. et al. Synthetic data in generalizable, learning-based neuroimaging. Imaging Neuroscience 2, imag–2 (2024).

[41] Hoffmann, M. Domain-randomized deep learning for neuroimage analysis: Selecting training strategies, navigating challenges, and maximizing benefits. IEEE signal processing magazine 42, 78–90 (2025).

[42] Billot, B. et al. SynthSeg: Segmentation of brain MRI scans of any contrast and resolution without retraining. Medical image analysis 86, 102789 (2023).

[43] Mauri, C. et al. A contrast-agnostic method for ultra-high resolution claustrum segmentation. Human Brain Mapping 46, e70303 (2025).

[44] Hoffmann, M. et al. SynthMorph: learning contrast-invariant registration without acquired images. IEEE transactions on medical imaging 41, 543–558 (2021).

[45] Walsh, C. L. et al. Imaging intact human organs with local resolution of cellular structures using hierarchical phase-contrast tomography. Nature methods 18, 1532–1541 (2021).

[46] Ronneberger, O., Fischer, P. & Brox, T. U-net: Convolutional networks for biomedical image segmentation. In Medical image computing and computer-assisted intervention–MICCAI 2015: 18th international conference, Munich, Germany, October 5-9, 2015, proceedings, part III 18, 234–241 (Springer, 2015).

[47] Dice, L. R. Measures of the amount of ecologic association between species. Ecology 26, 297–302 (1945).

[48] Duvernoy, H. M., Delon, S. & Vannson, J. Cortical blood vessels of the human brain. Brain research bulletin 7, 519–579 (1981).

[49] Taoka, T. et al. Structure of the medullary veins of the cerebral hemisphere and related disorders. Radiographics 37, 281–297 (2017).

[50] Costantini, I. et al. A cellular resolution atlas of broca’s area. Science Advances 9, eadg3844 (2023).

[51] Park, J. et al. Integrated platform for multiscale molecular imaging and phenotyping of the human brain. Science 384, eadh9979 (2024).

[52] Schilling, K. G. et al. The relationship of white matter tract orientation to vascular geometry in the human brain. Scientific Reports 15, 18396 (2025).

[53] Ogawa, S., Lee, T.-M., Kay, A. R. & Tank, D. W. Brain magnetic resonance imaging with contrast dependent on blood oxygenation. proceedings of the National Academy of Sciences 87, 9868–9872 (1990).

[54] Uludagğ, K., MuÜller-Bierl, B. & Ugğurbil, K. An integrative model for neuronal activity-induced signal changes for gradient and spin echo functional imaging. Neuroimage 48, 150–165 (2009).

[55] Kim, S.-G. & Ogawa, S. Biophysical and physiological origins of blood oxygenation level-dependent fmri signals. Journal of Cerebral Blood Flow & Metabolism 32, 1188–1206 (2012).

[56] Axer, M. & Amunts, K. Scale matters: The nested human connectome. Science 378, 500–504 (2022).

[57] Walsh, C. L. et al. The human organ atlas. Science Advances 12, eadz2240 (2026).

[58] Magnain, C. et al. Blockface histology with optical coherence tomography: a comparison with nissl staining. NeuroImage 84, 524–533 (2014).

[59] Liu, C. J. et al. Refractive-index matching enhanced polarization sensitive optical coherence tomography quantification in human brain tissue. Biomedical Optics Express 13, 358–372 (2021).

[60] Wang, H. et al. as-psoct: Volumetric microscopic imaging of human brain architecture and connectivity. NeuroImage 165, 56–68 (2018).

[61] Fischl, B. FreeSurfer. Neuroimage 62, 774–781 (2012).

[62] Manjoón, J. V., Coupeé, P., Martí-Bonmatí, L., Collins, D. L. & Robles, M. Adaptive non-local means denoising of mr images with spatially varying noise levels. Journal of Magnetic Resonance Imaging 31, 192–203 (2010).

